# Modulating pathological oscillations by rhythmic non-invasive brain stimulation – a therapeutic concept?

**DOI:** 10.1101/014548

**Authors:** LK Krawinkel, AK Engel, FC Hummel

**Author notes:** Correspondence: Friedhelm Hummel, BrainImaging and NeuroStimulation (BINS) Laboratory, Department of Neurology, University Medical Center Hamburg-Eppendorf, 20246 Hamburg, Germany.

## Abstract

A large amount of studies of the last decades revealed an association between human behaviour and oscillatory activity in the human brain. Alike, abnormalities of oscillatory activity were related with pathological behaviour in many neuropsychiatric disorders, such as in Parkinson’s disease (PD) or in schizophrenia (SCZ). As a therapeutic tool, non-invasive brain stimulation (NIBS) has demonstrated the potential to improve behavioural performance in patients suffering from neuropsychiatric disorders. Since evidence accumulates that NIBS might be able to modulate oscillatory activity and related behaviour in a scientific setting, this review focuses on discussing potential interventional strategies to target abnormalities in oscillatory activity in neuropsychiatric disorders. In particular, we will review oscillatory changes described in patients after stroke, with PD or suffering from SCZ. Potential ways of targeting interventionally the underlying pathological oscillations to improve related pathological behaviour will be further discussed.

## 1. Importance of oscillations for information processing in the healthy brain

Since Berger first described oscillatory activity of the brain in the 1920s, oscillations have been investigated extensively and research revealed clear relevance of human brain oscillations for information processing and behavior (Singer and Gray 1995; Engel et al. 2001; Brown 2003; Buzsáki and Draguhn 2004; Pfurtscheller et al. 2005; Schoffelen et al. 2005; Fries 2009; Jensen and Mazaheri 2010).

Generally, an oscillation is a periodic fluctuation caused by changes in excitability of a group of neurons, measured on the scalp as differences in voltage with techniques like electroencephalography (EEG) or magnetencephalography (MEG). Oscillations are suggested to be able to temporally coordinate and control neuronal firing and are proposed to be one basic principle of information processing in the human brain (Engel et al. 2001; Varela et al. 2001; Buzsáki and Draguhn 2004). Based on phenomenological observations, a division in frequency bands has been established. Usually, oscillation frequencies are categorized as delta-band (1.5–4 Hz), theta-band (4-7 Hz), alpha-band (8-12 Hz), beta-band (12-30 Hz) and gamma-band (> 30 Hz).

Slow delta waves are prominent in sleep and are, like theta-band oscillations, associated with memory consolidation (Marshall et al. 2006; Rasch and Born 2013). Alpha-band oscillations are the prominent oscillators in the resting brain and have been proposed to reflect functional inhibition (Klimesch et al. 2007; Jensen and Mazaheri 2010). In the motor system alpha-band oscillations decrease during performing a task and increase at rest (Pfurtscheller 1992) or during inhibitory control processes (Hummel et al. 2002; Sauseng et al. 2013). In line with these results, task-related alpha activity decreases in cortical regions involved in the task but increases in regions that are not involved as a correlate of deactivation. These results led to the hypothesis that alpha-band activity reflects inhibition (Haegens et al. 2010). Beta-band oscillations are, like alpha oscillations, associated with motor control (Kilner et al. 2000; Pfurtscheller et al. 2005). Since beta oscillations are described to be associated with a steady state of the motor system, it has been hypothesised that they signal the ‘status quo’ (Gilbertson et al. 2005; Engel and Fries 2010). Having been extensively investigated, gamma-band oscillations are thought to be involved into a broad range of behavioural components like visual perception and attention (Singer and Gray 1995; Jensen et al. 2007) and were associated with memory functions (Tallon-Baudry et al. 1998; Herrmann et al. 2004).

Moreover, mechanisms of oscillatory communication have been identified. For example synchronisation of phases is suggested to be one basic principle for long range communication in the brain (Varela et al. 2001; Hummel and Gerloff 2005; Schoffelen et al. 2005). Furthermore high frequency activity has been described as being depended on low frequency phase (Osipova et al. 2008; Jensen and Mazaheri 2010; Buzsáki and Wang 2012).

Underlining a potential causal relationship, both oscillatory activity and behaviour were modulated by NIBS in several studies. Before focussing on abnormal oscillations and pathological behavior, the influence of NIBS on oscillations of the healthy brain will be discussed in the next section.

## 2. Modulation of oscillations by rhythmic NIBS in the healthy brain

Several recent studies combining non-invasive brain stimulation and EEG revealed that NIBS is able to modulate oscillations in the healthy brain (Plewnia et al. 2008; Pogosyan et al. 2009; Zaehle et al. 2010; Thut et al. 2011; Vernet et al. 2013). Generally, one basic technique of NIBS is Transcranial Magnetic Stimulation (TMS). By depolarization of neurons, a TMS single pulse applied over the hand knob area typically leads to a motor evoked potential in the contralateral hand (Hummel and Cohen 2005; Hallett 2007). Combining TMS with EEG, an increase of the intrinsic frequency of the TMS-affected area and their thalamic loops respectively has been observed (Rosanova et al. 2009). Another basic method of NIBS is Transcranial Electric Stimulation (TES). In contrast to TMS, TES is suggested to tune oscillators by modulating the membrane potential of neurons and their spontaneous firing rates (Nitsche and Paulus 2000).

Regarding rhythmic applications of TMS, there is e.g., the theta burst stimulation (TBS) protocol, in which a triple pulse with an interpulse frequency of 50 Hz is applied in a theta rhythm. Applied continuously, TBS has an inhibitory effect, applied in an intermittent mode TBS has an excitatory effect (Huang et al. 2005; Hallett 2007). Furthermore, repetitive TMS applied in a frequency below or at 1 Hz has an inhibitory effect, application of frequencies at faster rates (>5 Hz) leads to excitation (Pascual-Leone et al. 1994; Hummel and Cohen 2005). Regarding patterned applications of TES, there is transcranial alternating current stimulation (tACS), a sinusoidal stimulation with spatially alternating anodal and cathodal components. Furthermore, there are transcranial random noise stimulation (tRNS) and direct current stimulation with a sinusoidal component, called oscillatory tDCS (Siebner and Ziemann 2010).

Depending on their effect on oscillations, a division of rhythmic applications of NIBS into two concepts has been proposed (Siebner and Ziemann 2010). On the one hand rhythmic NIBS can be applied in an oscillatory mode in the frequency of the targeted oscillation or its oscillator, respectively (Pogosyan et al. 2009; Zaehle et al. 2010; Thut et al. 2011). In the following, this mode will be called ‘direct’, because there is a direct interference of stimulation frequency and the frequency of the oscillator. On the other hand, one can apply rhythmic NIBS in a non oscillatory mode, not reflecting the targeted oscillations or the frequency of the targeted oscillator. In this case NIBS is suggested to modulate underlying oscillatory mechanisms (Vernet et al. 2013; Nitsche and Paulus 2000; Lapenta et al. 2013). This non oscillatory mode will be called ‘indirect’ in the following.

In a study on ‘direct’ modulation of ongoing oscillations, five pulses of TMS with an adjusted individual alpha frequency were able to entrain a parietal alpha oscillator specifically in its natural frequency (Thut et al. 2011). tACS entrained individual parietal-central alpha oscillations if applied in the endogenous frequency at the occipital pole (Zaehle et al. 2010). Furthermore, 10 Hz repetitive bifocal TMS over left primary motor cortex and over visual cortex led to an increase of alpha-band and lower beta coherence between the stimulated sites (Plewnia et al. 2008), supporting the exciting concept to not only influence local oscillatory activity but also long-range oscillatory interactions by NIBS.

Combining NIBS with EEG, ‘direct’ applications of NIBS also modulated both oscillatory activity and behavioral components, thereby providing evidence for a causal role of oscillations and its modulation. In a visuomotor task tACS applied at 20 Hz reduced peak velocity on the one hand and increased coherence between scalp-recorded activity and EMG activity at 20 Hz on the other hand (Pogosyan et al. 2009). Recently it has been shown by concurrent EEG-tACS that 10 Hz stimulation can entrain parieto-occipital alpha activity and modulate target detection performance in an oddball task in a phase-dependent manner (Helfrich et al. 2014). In an auditory experiment perception thresholds were dependent on the phase of the entrained oscillation using 10 Hz oscillatory tDCS (Neuling et al. 2012). Furthermore slow wave oscillatory tDCS in a frequency of 0.75 Hertz applied during non-rapid-eye-movement sleep induced an increase of slow wave oscillations and enhanced the retention of declarative memories (Marshall et al. 2006). However, this study was critically evaluated by the same group showing that the total amount of current and not the oscillatory component of the oscillatory tDCS might have been the main effective variable (Groppa et al. 2010).

As an ‘indirect’ rhythmic application of NIBS, continuous theta burst stimulation (cTBS) increased theta-band power and decreased beta-band power in an eyes-closed resting experiment (Vernet et al. 2013). As an underlying mechanism the authors suggest that cTBS modulates synchronisation of relevant oscillators. In a non resting setup cTBS increased event-related lower beta power applied over primary motor cortex for at least 30 minutes (Noh et al. 2012) and 40 trains of ten excitatory TMS pulses at 20 Hz increased alpha and beta-band event-related synchronization at the stimulation site (Veniero et al. 2011).

Like ‘direct’ applications, also ‘indirect’ applications were able to modulate both oscillations and behaviour. Inhibitory 1 Hz repetitive TMS on right prefrontal cortex reduced both response times for congruent cued targets and ipsilateral alpha amplitude (Sauseng et al. 2011). In another study 5 Hz tDCS (theta tDCS) decreased both slow wave activity, frontal slow EEG spindle power and consolidation of declarative memory (Marshall et al. 2011).

Taken together evidence is increasing that oscillations, being one basic principle of information processing in the human brain, can be modulated by NIBS. In many cases, this suggests a possible mechanism for how NIBS may exert its effects on cognitive processes or behaviour.

In the following paragraph, alteration of oscillatory activity will be discussed as a pathophysiological mechanism in neuropsychiatric disorders. As examples, we will consider oscillatory changes in the alpha-band after stroke, beta-band changes in patients with Parkinson’s disease (PD) and altered gamma-band activity in patients suffering from schizophrenia (SCZ). Next, we will discuss potential applications of rhythmic NIBS like repetitive TMS and tACS in order to modulate pathological oscillations and potentially improve the clinical outcome in patients.

## 3. Role of oscillatory activity in the pathophysiology of neuropsychiatric disorders

Stroke is one of the leading causes for acquired long-term disability in industrialized countries (Kolominsky-Rabas et al. 2001) and, therefore, studies on neurophysiological changes accompanying and following stroke have received considerable interest.

Several studies have observed changes in alpha-band activity recorded over the affected hemisphere (AH) after stroke in the resting brain (Tecchio et al. 2005, 2006; Dubovik et al. 2012; Westlake et al. 2012; Laaksonen et al. 2013). 32 patients with stroke of the middle cerebral artery within the first 10 days (Tecchio et al. 2005) and 56 stroke patients in a chronic stage (Tecchio et al. 2006) showed a reduction of the individual alpha frequency in the AH. In 16 patients with affected upper limp function, amplitude of alpha-band oscillations increased in the AH compared to the unaffected hemisphere (UH) and to control subjects one month and three months after stroke (Laaksonen et al. 2013). In the latter study, alpha oscillations had a burst-like pattern and were found both in rolandic and in parietal regions. However, these changes in alpha-band activity did not correlate with the clinical outcome in patients (Tecchio et al. 2005, 2006; Laaksonen et al. 2013).

In contrast, two studies addressing interregional resting-state functional connectivity (FC) were able to correlate changes of alpha-band activity with performance (Dubovik et al. 2012) and motor recovery (Westlake et al. 2012) after stroke. 20 patients showed a decrease of alpha-band FC of central electrodes over the lesions to all other electrodes three months after stroke of middle or/and anterior cerebral artery (Dubovik et al. 2012). Moreover, motor functions correlated with IC values of the precentral gyrus and all other investigated brain regions. Since FC changes were not restricted to the boundaries of the lesions but restricted to a specific frequency, the authors emphasize that oscillatory changes are probably not due to tissue loss but to changes in affected tissue. Furthermore the study showed that changes in FC are probably due to changes in alpha phase synchrony rather than to changes in alpha amplitude (Stam et al. 2007; Dubovik et al. 2012). Another study found changes in alpha-band connectivity in the acute phase of stroke (Westlake et al. 2012). Greater initial functional alpha-band connectivity of ipsilesional primary sensory cortex and prefrontal cortex in the acute phase correlated with better clinical improvement 8 two 12 weeks after stroke in fourteen patients with motor impairment of the upper limb.

Next to changes in alpha-band activity, also changes of slow wave oscillations were found after brain injuries like head trauma and stroke (Lewine et al. 1999; Butz et al. 2004). In line with these results a shift from fast to slow rhythms could be observed after stroke (Dubovik et al. 2012) and slow delta wave oscillations of the unaffected hemisphere have been shown to be correlated with clinical outcome after stroke (Tecchio et al. 2007). Patients with persisting abnormal slow wave oscillations (abnormal low frequency magnetic activity) had a significantly worse clinical outcome compared to patients without persisting slow wave oscillatory components (Laaksonen et al. 2013).

Besides clinical observations, slow wave activity has also been investigated in an animal stroke model, in which slow wave oscillations were associated with axonal sprouting after thermal-ischemic lesioning (Carmichael and Chesselet 2002). In this model, a treatment with tetrodotoxin reduced both slow oscillations and axonal sprouting. In another study neuronal bursts were able to reduce inhibitory factors that surround axons (Ming et al. 2001). In a recent publication it could be shown that new patterns of axonal sprouting mediated recovery after stroke in an animal model (Overman et al. 2012). Taken together, these data provided first evidence that there might be an association of slow neuronal firing, axonal sprouting and behavioral recovery.

Parkinson’s disease has a high prevalence in aging populations (von Campenhausen et al. 2005). Evidence is increasing that clinical symptoms of PD are related to abnormalities in beta-band activity (Levy et al. 2002; Kühn et al. 2008, 2009; Crowell et al. 2012; Heinrichs-Graham et al. 2014; Herz et al. 2014). Prominent beta-band activity has been recorded in the basal ganglia in animal models of PD and in PD patients with deep brain stimulation (Levy et al. 2000, 2002; Kühn et al. 2008; Bergman et al. 2013; Heinrichs-Graham et al. 2014; Herz et al. 2014). Both dopamine treatment and DBS have been shown to reduce beta-band activity (Brown et al. 2001; Levy et al. 2002; Kühn et al. 2008). The modulation of beta-band activity by dopaminergic treatment (Kühn et al. 2009) and by DBS (Ray et al. 2008) was associated with a reduction of bradykinesia and rigidity. A recent study in PD also revealed that treatment with levodopa reinforced beta-band coupling between primary motor cortex and lateral premotor cortex (Herz et al. 2014). Another study demonstrated a lack of physiological event-related beta desynchronization which could be modulated by levodopa in PD (Heinrichs-Graham et al. 2014). Taken together, in PD abnormal beta oscillations are a consistent finding with a strong relation to abnormal behaviour. Hence, beta-band oscillations could be a promising target for interventional strategies based on rhythmic NIBS.

Schizophrenia has a life-time prevalence of 0.4% and belongs to the thirty most disabling disorders worldwide (Murray and Lopez 1997; McGrath et al. 2008). Besides psychotic symptoms patients with SCZ show abnormalities in cognitive performance, e.g., in working memory, cognitive control and sensory gating. These cognitive deficits are, on the one hand, associated with clinical and functional outcome (Green et al. 2000; Gold 2004; McGrath et al. 2008). On the other hand they are related with abnormalities in gamma-band oscillations (Johannesen et al. 2005; Uhlhaas et al. 2008; Barr et al. 2010; Sun et al. 2011). Compared to healthy controls, patients with SCZ showed excessive gamma-band oscillations in conditions with high memory load (Barr et al. 2010). Altered gamma-band connectivity was also found in the resting brain of SCZ patients (Andreou et al. 2014). Furthermore, evoked gamma power and phase synchronization were reduced after auditory stimulation in patients with SCZ (Light et al. 2006). In line with these results, after a 40 Hz binaural stimulation a reduced phase locking between primary auditory cortices was observed in patients with SCZ, which correlated with auditory hallucination scores (Mulert et al. 2011).

Moreover, combination of TMS and EEG has been used in SCZ research by using a TMS double pulse paradigm (Farzan et al. 2009) called long intracortical inhibition (LICI). LICI is an established, probably GABA_B_ mediated measurement for cortical inhibition in motor cortex using motor-evoked potentials as a readout (McDonnell et al. 2006). Both GABA_A_ and GABA_B_ suggested to be involved in the generation of gamma-band oscillations (Wang and Buzsáki 1996; Brown et al. 2007). Recently, EEG parameters were established as another readout for double pulse TMS paradigms in non motor areas like DLPFC (Daskalakis et al. 2008). A study in healthy subjects demonstrated that applying a TMS LICI protocol over DLPFC led to a decrease of gamma oscillations whereas gamma oscillations over motor cortex did not change after LICI application (Farzan et al. 2009). In contrast, patients with SCZ showed an impaired LICI-mediated inhibition of gamma oscillations in DLPFC compared to healthy participants and patients with bipolar disorder (Farzan et al. 2010a).

In summary, gamma-band oscillations have been related with cognitive performance in the healthy brain (Tallon-Baudry et al. 1998; Herrmann et al. 2004). In SCZ abnormal gamma-band activity was associated with behavioural impairment (Johannesen et al. 2005; Uhlhaas et al. 2008; Barr et al. 2010; Sun et al. 2011; Andreou et al. 2014). In line with these results, gamma-band activity could be one potential target of NIBS in SCZ.

## 4. Targeting pathological oscillations by non-invasive brain stimulation

This section will address the question of how rhythmic NIBS could be used to modulate pathological oscillations and thereby modulate behavior and clinical symptoms in stroke, PD and SCZ.

Several studies have demonstrated that NIBS can improve functional outcome of stroke patients (Hummel and Cohen 2005; Hummel et al. 2005, 2006; Kim et al. 2006; Cazzoli et al. 2012; for review Schulz et al. 2013). To our knowledge, however, there is to date limited data on the effect of NIBS on oscillations in stroke. In six patients with severe hemiparesis, tDCS modulated alpha-band oscillations of sensorimotor areas during imaginary movements of the affected hand (Kasashima et al. 2012). Since detection of stable brain rhythms can be difficult in damaged brain areas after stroke, the aim of the latter study was to enhance oscillatory activity in the lesioned hemisphere by tDCS in order to be able to achieve a stable and well distinguishable brain signal for use in a brain-computer-interface (BCI). Event related alpha-band desynchronisation, a correlate of local activation, was significantly enhanced after tDCS stimulation compared to sham stimulation, making the signal more suitable for a BCI.

Although several studies have applied NIBS in stroke patients, only one case report has used rhythmic NIBS to modulate oscillations. In a patient with aphasia 10 Hz rTMS modulated both oscillatory activity and clinical outcome (Dammekens et al. 2014). In the treated patient the stroke lesion affected the left inferior frontal gyrus (IFG), which is part of the language network (Crosson et al. 2007). Generally, according to the concept of an interhemispheric rivalry in stroke (Murase et al. 2004; Duque et al. 2005), inhibitory rTMS protocols of 1 Hz stimulation frequency are applied to the healthy hemisphere in order to prevent interhemispheric maladaptive processes in contralesional networks. Another approach is to excite the damaged hemisphere with rTMS protocols with a frequency above 5 Hz (Lefaucheur 2006). In the latter study, excitatory rTMS was applied over the damaged left IFG in daily sessions over three weeks leading to an increase of functional connectivity between left and right IFG in the theta-and high beta-band. Behaviourally, the patient improved on repetition tasks for naming and comprehension. In addition to being a case report, this study does certainly not establish a causal relation between modulation of oscillations and changes in behavior. Nevertheless, this study provides first hints that rhythmic NIBS might be effective in modulating both functional connectivity and clinical outcome.

Based on the emerging understanding about the relationship between NIBS and the modulation or induction of oscillatory activity in the last decad, hypotheses on how to enhance brain functioning and ameliorate impaired behaviour after focal brain lesions can be defined. These hypotheses would be amenable to evaluation in controlled clinical trials in stroke patients. Combining the evidence that (1) changes in alpha-band activity are correlated with recovery and performance after stroke (Dubovik et al. 2012; Westlake et al. 2012) and that (2) NIBS is able modulate alpha-band activity (Plewnia et al. 2008; Thut et al. 2011) one could consider applying rhythmic alpha-NIBS after stroke.

As illustrated schematically in Figure 1A, bifocal TMS at alpha frequency could enhance functional connectivity between lesioned areas and relevant neighbouring regions. In this context, bifocal application should potentially be performed with a certain phase lag between the targeted areas. An application of less focal alpha tACS in a montage covering both targeted areas would be another option. In both cases rhythmic NIBS would potentially lead to a synchronization in the alpha-band of lesioned sites and relevant connected areas like ipsilateral and contralateral premotor cortices respectively (Johansen-Berg et al. 2002; Ward et al. 2003; Gerloff et al. 2006; Rehme et al. 2011; Dubovik et al. 2012; Westlake et al. 2012).

**Figure 1A.**
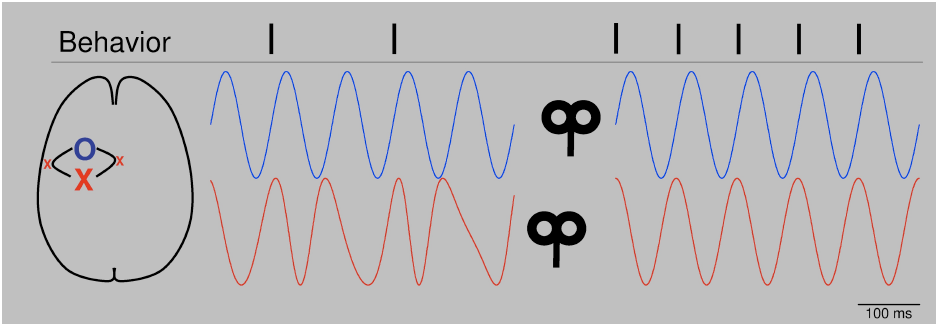
Schematic: Bifocal repetitive TMS restores functional connectivity. After stroke, alpha oscillators (red line, left) of the affected area (red X, primary motor cortex) show unsteady phase properties. Since physiological behavior depends on a constant phase relation of alpha oscillations between the affected area and connected networks (blue O, blue line, left, premotor areas), the patient is impaired (behavior, left, e.g., muscle contractions). After application of bifocal, repetitive TMS at alpha frequency over both areas with a lag reflecting physiological offset of oscillators, oscillatory activity is synchronised (blue and red line, right) in terms of a constant phase lag. Behavior improves (right). Please note that conditions are highly simplified. First, next to phase, also amplitude is important for signal processing. Second, in the presented physiological condition (right) phases are completely locked, which is artificial. Naturally, phases would fluctuate.

Another option to increase alpha-band connectivity could be to add noise, as shown in Figure 1B. Transcranial random noise stimulation (tRNS) is a technique which has been used to modulate cortical excitability and BOLD activity (Terney et al. 2008; Saiote et al. 2013). It has been suggested that, by adding noise, tRNS is able to augment cortical oscillations of different frequencies (Antal and Paulus 2013). In stroke patients, tRNS could elevate reduced alpha connectivity in affected parts of the brain back to suprathreshold levels (Moss et al. 2004) and thereby restore functional connectivity.

**Figure 1B.**
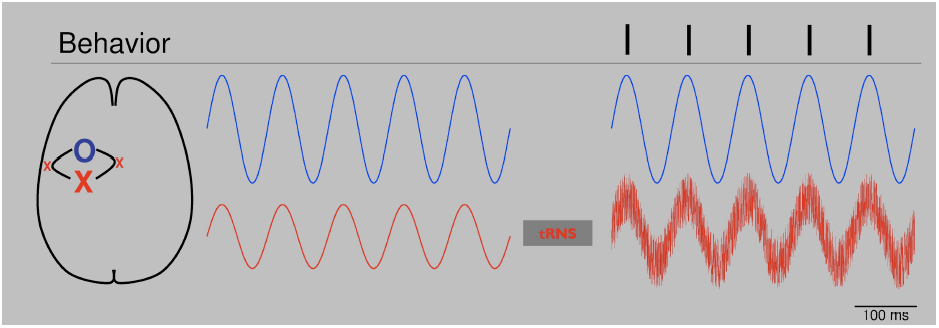
Schematic: tRNS augments alpha oscillators. After stroke alpha activity (red line, left) of the affected area (red X) is reduced. Threshold for establishing information processing with a related network (blue line, left) is not reached and dependent normal behavior is reduced. tRNS applied over the affected oscillator augments alpha activity by adding noise (red line, right). Information processing is re-established and behavior improves. Please note that conditions are very much simplified. In both cases, phase is totally locked without any phase lag. Biological signals would fluctuate.

Regarding the results of reduced alpha peak frequency and increased alpha amplitude in the affected hemisphere after stroke (Tecchio et al. 2005) one could argue that behavioral changes might be related to alpha induced inhibition (Jensen and Mazaheri 2010). In contrast to changes in functional connectivity, these changes were not correlated with behavioural changes, however, and gaining clinical benefit by modulating these features of local oscillatory activity remains an untested possibility.

In addition to the modulation of electrophysiological dynamics, NIBS might also influence molecular environment of neurons. By modulating slow wave activity occurring after stroke (Tecchio et al. 2005; Dubovik et al. 2012; Laaksonen et al. 2013) related neural repair might be modulated. Since there is first evidence in an animal model that there might be an association of slow neuronal firing, axonal recovery and recovery of function (Ming et al. 2001; Carmichael and Chesselet 2002; Overman et al. 2012), one should consider a slow frequency stimulation as performed before in different contexts (Marshall et al. 2011) to gain axonal sprouting and improve clinical outcome after brain injury like stroke.

As discussed in the preceding section, pathological beta oscillations are a consistent finding in basal ganglia correlated with behavioural changes in PD patients (Levy et al. 2002; Kühn et al. 2008, 2009; Crowell et al. 2012; Heinrichs-Graham et al. 2014; Herz et al. 2014). It is known that these abnormal beta-band oscillations can be modulated by established deep brain stimulation (DBS) protocols. In addition, a number of studies investigated the effects of NIBS on motor function in patients with PD. Applying 5 Hz repetitive TMS (rTMS) over primary motor cortex led to a short term increase of motor function in six PD patients (Siebner et al. 2000). A 5 Hz rTMS protocol consisting of 2000 pulses per day over ten days led to significant reduction of motor impairment (Khedr et al. 2003). Moreover, rTMS over inferior frontal cortex modulated event-related potentials in the subthalamic nucleus (STN) in patients with implanted DBS electrodes (Rektor et al. 2010).

In our view, both the clinical effectiveness of DBS and of NIBS in modulating cortex-basal ganglia dynamics point to the potential of NIBS applications in PD. As discussed above, beta oscillations could be a promising target. A recent study has demonstrated the possibility to enhance the ‘akinetic’ effect of beta oscillations. In a task where subjects got a cue to stop a planned grip-movement, tACS stimulation with 20 Hz significantly reduced grip force in that condition (Joundi et al. 2012). One the one hand these results underline the potential impact of enhanced beta oscillations in ‘akinetic’ disorders like Parkinson’s disease. On the other hand, the results point out the ability of cortical NIBS to affect loops involving the basal ganglia.

To our knowledge, however, it has not been possible to reduce pathological beta oscillations in PD by NIBS. Potentially, reducing beta activity could be achieved by phase cancellation. Phase cancellation by application of tACS has already been used for tremor reduction in PD. Targeting tremor-associated oscillations in such an approach in patients with tremor-dominant PD led to an almost 50% suppression of resting tremor (Brittain et al. 2013). As a first step in the study, the individual tremor frequency of each patient was measured. In a second step, tremor-frequency adjusted tACS was applied over the primary motor cortex at different phase angles. Certain phase angles led to tremor reduction whereas stimulation at different angles led to an increase of tremor. This study underlines the great potential of individualized application of rhythmic NIBS and suggests the potential efficacy of phase cancellation. Based on monkey data, phase cancellation has also been suggested as a physiological mechanism for controlling tremor in spinal cord networks (Williams et al. 2010). As illustrated schematically in Figure 2A, phase cancellation might be applied to pathological beta oscillations in PD potentially targeting bradykinesia and rigidity (Ray et al. 2008; Kühn et al. 2009).

**Figure 2A.**
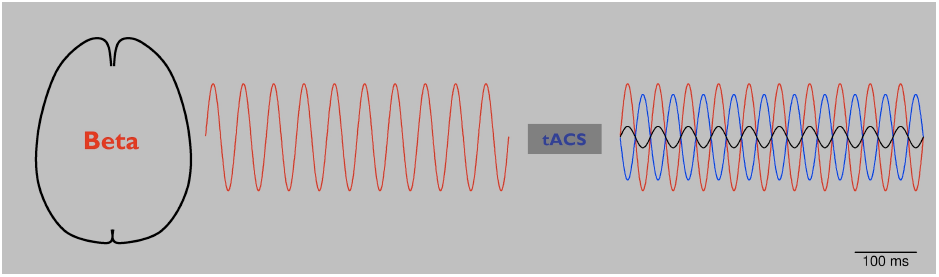
Schematic: tACS in Parkinson’s disease (PD). Abnormal beta oscillations (red line, left and right) are related to rigidity and bradykinesia. Application of tACS with a phase lag of 180° (blue line, right) leads to a reduction of these pathological oscillations (black line, right) by phase cancellation (right). As a result, rigidity and bradykinesia are reduced. Please note that to date phase cancellation has been thought to be able to reduce tremor in PD by affecting tremor related cortical frequencies. To our knowledge, beta frequency has not yet been targeted by NIBS using the phase cancellation concept.

Since tremor shifted to a different frequency in some of the patients in the study by Brittain and coworkers (2013), tACS application should be adjusted to ongoing tremor activity to provide an effective modulation, as shown schematically in Figure 2B. To this end, online read-out techniques that allow stimulation in dependence on ongoing oscillatory activity (Berényi et al. 2012; Bergmann et al. 2012) will be highly advantageous in order to adjust NIBS to changing pathologic frequency and provide effective phase cancellation and reduction of symptoms. In a clinical setting such an ‘online’, closed-loop approach has already been tested successfully: in PD DBS has been triggered based on recorded local field potentials (Little et al. 2013). These interventional strategies are interesting and potentially promising for amelioration of symptoms not well responsive to DBS.

**Figure 2B.**
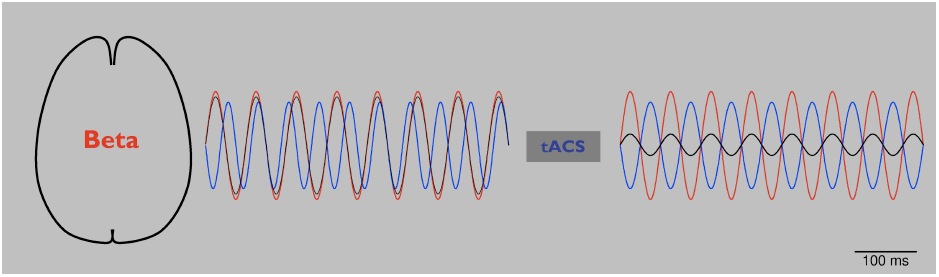
Schematic: tACS in Parkinson’s disease (PD) using online readout techniques. Since it has been reported that during tACS application tremor frequency changed in patients with PD, more online readout techniques should be developed to adjust NIBS applications. After application of 20 Hz tACS (see figure 2A), pathological beta rhythm changes its frequency to 15 Hz (red line, left). tACS of 20 Hz with 180° phase lag (blue line, left) looses its effect (black line, left). After online adjustment, tACS shifts to 15 Hz (blue line, right). Pathological beta (red line, right) is again reduced (black line, right).

In SCZ, combining single or double pulse TMS with EEG revealed abnormalities of cortical inhibition of gamma oscillations (Farzan et al. 2009, 2010b). Both studies point towards a scientific benefit of combining these methods and suggest that measurements on TMS-evoked oscillatory activity could be useful as a ‘biomarker’ for neuropsychiatric disorders. Applying a similar approach, TMS evoked EEG activity has been proposed to add information to diagnostic procedures for chronic disorders of consciousness (Ragazzoni et al. 2013).

Abnormal gamma-band oscillations of patients with SCZ have been modulated by repetitive TMS (Barr et al. 2011). In an N-back working memory task, subjects were asked to compare stimuli with those from previous trials. 20 Hz TMS applied bilaterally to DLPFC in reduced gamma activity in the SCZ patients (Barr et al. 2011). This approach, which is schematically presented in Figure 2C, could be considered as an ‘indirect’ application, in which underlying gamma oscillators were modulated by an excitatory 20 Hz rTMS protocol not directly targeted at gamma frequencies. Since repetitive TMS of a single session in the latter study did not change working memory behaviour, the same group performed a pilot study with 20 Hz rTMS over DLPFC bilateraly for twenty sessions in a randomized double-blinded sham-controlled manner. This application led to a significant improvement of accuracy in working memory underlining that repetitive TMS might potentially improve cognitive impairments in SCZ patients (Barr et al. 2013). Moreover, these results point towards the need for repetitive sessions of NIBS to gain long term effects. However, changes in gamma oscillations were not addressed in this follow-up study (Barr et al. 2013).

**Figure 2C.**
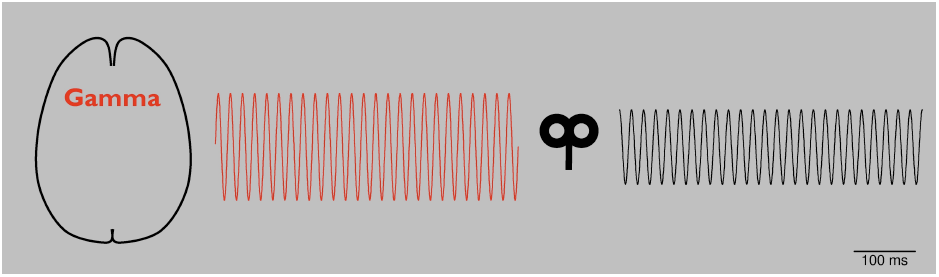
Schematic: Repetitive TMS in schizophrenia (SCZ). By modulating oscillators of abnormal gamma-band activity (red line, left), amplitude is reduced (black line, right) and related behavioral impairments are reduced.

Taken together, application of rTMS seem to be promising for the treatment of negative symptoms of SCZ, although so far only demonstrated in proof-of-principle studies. Gamma oscillations show a high diversity in SCZ (Sun et al. 2011) and, hence, a direct link between the modulation of gamma oscillations and an improved cognitive performance in SCZ remains speculative. Nevertheless, further applications for the treatment of negative symptoms based on safe and low cost NIBS techniques addressing specifically gamma oscillations should be further developed and evaluated in detail in future studies.

## 5. Summary and further perspectives of NIBS

Evidence is increasing that NIBS may provide a novel and promising strategy to modulate both oscillatory activity and related behavior in the healthy brain. To date only a limited number of studies have employed NIBS to modulate oscillatory abnormalities in neuropsychiatric disorders as a treatment strategy. As discussed above, this concept is supported by studies suggesting, for example, that tACS can reduce tremor in PD (Brittain et al. 2013) or that rhythmic TMS can improve both working memory (Barr et al. 2013) and impaired gamma activity in patients with SCZ (Barr et al. 2011). In stroke, NIBS might modulate both oscillatory activity and clinical performance (Dammekens et al. 2014). Taken together, the application of rhythmic NIBS in order to modulate underlying, disease related oscillations is a very promising approach.

Next steps will be the development and evaluation of safe and low-cost applications of patterned NIBS in order to target pathological oscillatory activity to achieve improvement of clinical symptoms of neuropsychiatric patients impacting on their daily life. Demonstrating the expected effects of NIBS in patients might have an incremental impact on treatment of neuropsychiatric disorders and subsequently the health system, as novel devices has been developed for home-based, self-application of NIBS. As soon as such approaches will be proven save in the home-based environment, they may offer a cost-effective strategy to significantly enhance treatment intensity.

Still one can argue critically that oscillatory changes after NIBS application are not a cause but a byproduct merely associated with behavioral changes. However, combining NIBS with electrophysiological recordings bears great potential to establish the specificity and reliability of oscillations as biomarkers.

## 6. Acknowledgement

This work has been supported by the German Research Foundation (DFG, SFB936/A2/A3/C4) and by the European Union (ERC-2010-AdG-269716 „MULTISENSE“).

